# Neural codes of seeing architectural styles

**DOI:** 10.1101/045245

**Authors:** Heeyoung Choo, Jack Nasar, Bardia Nikrahei, Dirk B. Walther

## Abstract

Images of iconic buildings, such as the CN Tower, instantly transport us to specific places, such as Toronto. Despite the substantial impact of architectural design on people’s visual experience of built environments, we know little about its neural representation in the human brain. In the present study, we have found patterns of neural activity associated with specific architectural styles in several high-level visual brain regions, but not in primary visual cortex (V1). This finding suggests that the neural correlates of the visual perception of architectural styles stem from style-specific complex visual structure beyond the simple features computed in V1. Surprisingly, the network of brain regions representing architectural styles included the fusiform face area (FFA) in addition to several scene-selective regions. Hierarchical clustering of error patterns further revealed that the FFA participated to a much larger extent in the neural encoding of architectural styles than entry-level scene categories. We conclude that the FFA is involved in fine-grained neural encoding of scenes at a subordinate-level, in our case, architectural styles of buildings. This study for the first time shows how the human visual system encodes visual aspects of architecture, one of the predominant and longest-lasting artefacts of human culture.

---

As of 2014, more than half of the world’s population resided in urban environments [1]. Architectural design has profound impact on people’s preferences and productivity in such built environments [2, 3]. Despite the ubiquity and importance of architecture for people’s lives, it is so far unknown where and how architectural styles are represented in people’s brains. While appraisal of architectural design is a collective experience encompassing perceptual, cognitive, and emotional experiences, architecture takes essentially a visual form [2]. That is, even though people have different cognitive interpretations and emotional responses to the Walt Disney Concert Hall in Los Angeles, they are likely to agree that the building exhibits an unusual asymmetric shape composed of metallic exterior surfaces with high curvature (shown in Fig 1C. Gehry). Here we show that the perceptual basis of architectural styles is represented in distributed patterns of neural activity in several visually active brain regions in ventral temporal cortex, but not in primary visual cortex.

**Figure 1.**
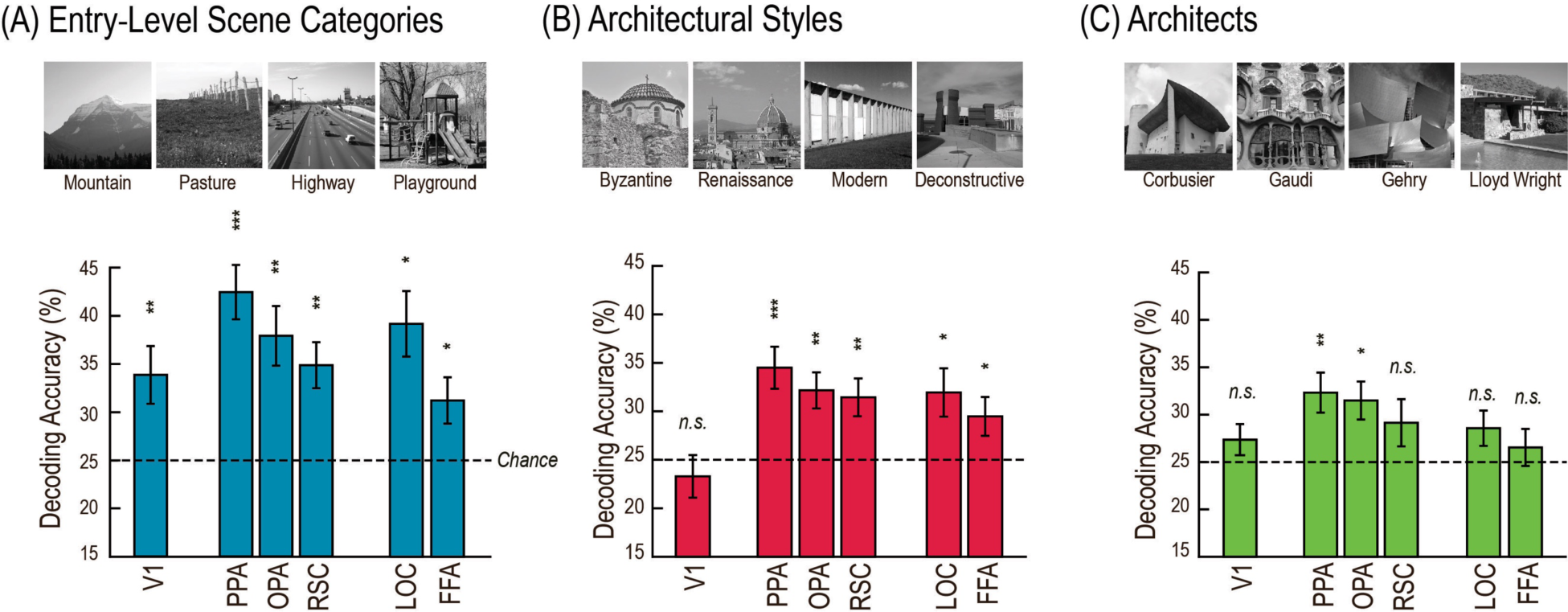
Example images and category decoding accuracy rates for three visual categories across the ROIs: (A) entry-level scene categories, (B) architectural style, and (C) architects (for differences in mean activity levels see Fig S1). These public-domain example images were not shown to the participants, but are visually similar to the experiment stimuli (i.e., depicting the same architecture) downloaded from the World Wide Web. Decoding of face identity was only possible in V1 (at 37.1%, *p* = 6.18·10^−5^) and is not shown here. Error bars indicate standard errors of mean. Significance with respect to chance (25%) was assessed at the group level with one-sample t-tests (one-tailed). P-values were adjusted using false discovery rate, \**p* < .05, \*\**p* .01, \*\*\**p* < .001.

In a functional magnetic resonance imaging (fMRI) scanner, 23 students in their final year at The Ohio State University (11 majoring in architecture, 12 majoring in psychology or neuroscience, one psychology student excluded due to excessive head motion) viewed blocks of images while performing a one-back task. Each block comprised four images from one of the following sixteen categories; (1) representative buildings of four architectural styles (Byzantine, Renaissance, Modern, and Deconstructive); (2) representative buildings designed by four famous architects of Modern and Deconstructive styles (Le Corbusier, Antoni Gaudi, Frank Gehry, and Frank Lloyd-Wright); (3) four entry-level scene categories (mountains, pastures, highways, and playgrounds); and (4) photographs of faces of four different non-famous men (Fig. 1). The building images encompassed a variety of views, including close-ups of signature facets of an architecture, far views capturing an entire building, and aerial views. Brain activity was recorded in 35 coronal slices, which covered approximately the posterior 70% of the brain. For each participant, several visually active regions of interest (ROI) were functionally localized: the parahippocampal place area (PPA), the occipital place area (OPA), the retrosplenial cortex (RSC), the lateral occipital complex (LOC), and the fusiform face area (FFA). Primary visual cortex (V1) was defined on each participant’s original cortical surface map using the automatic cortical parcellation provided by Freesurfer [4]. Surface-defined V1 was registered back to the volumetric brain separately for each hemisphere using AFNI (see Supplementary Methods for the ROI delineation methods).

Following standard pre-processing, data from the image blocks were subjected to a multi-voxel pattern analysis (MVPA). For each of the four groups of stimuli, a linear support vector machine decoder was trained to discriminate between the activity patterns associated with each of the four sub-categories. The decoder was tested on independent data in a leave-one-run-out (LORO) cross validation. Separate decoders were trained and tested for each participant and each ROI. Accuracy was compared to chance (25%) at the group level using one-tailed t tests.

## Results

### Successful decoding of architectural categories from human visual cortex

We used a mixed analysis of variance (ANOVA) to test for differences in decoding accuracy between experts and non-experts. We found no differences between the groups and therefore proceeded to collapse the data for participants from both groups for further analysis. The details of the inter-group analysis are discussed at the end of the results section.

Consistent with previous results [5, 6, 7], we could decode entry-level scene categories from all visually active ROIs (Fig. 2A). Furthermore, we could decode architectural styles from all five high-level visual brain regions, but not from V1 (Fig. 2B). In addition, it was possible to decode buildings by famous architects from brain activity in the PPA and the OPA, but not from V1, the RSC, the LOC, or the FFA (Fig. 2C). Decoding of facial identity succeeded only in V1 and was not possible in any of the high-level ROIs, including the FFA. Supplementary Table S1 shows details of the statistical results. Full discrimination between sub-categories was only possible by considering the *spatial patterns* of brain activity within ROIs (see Supplementary Figure S1 for mean neural activity results, and Supplementary Table S2 for univariate LORO decoding results).

**Figure 2.**
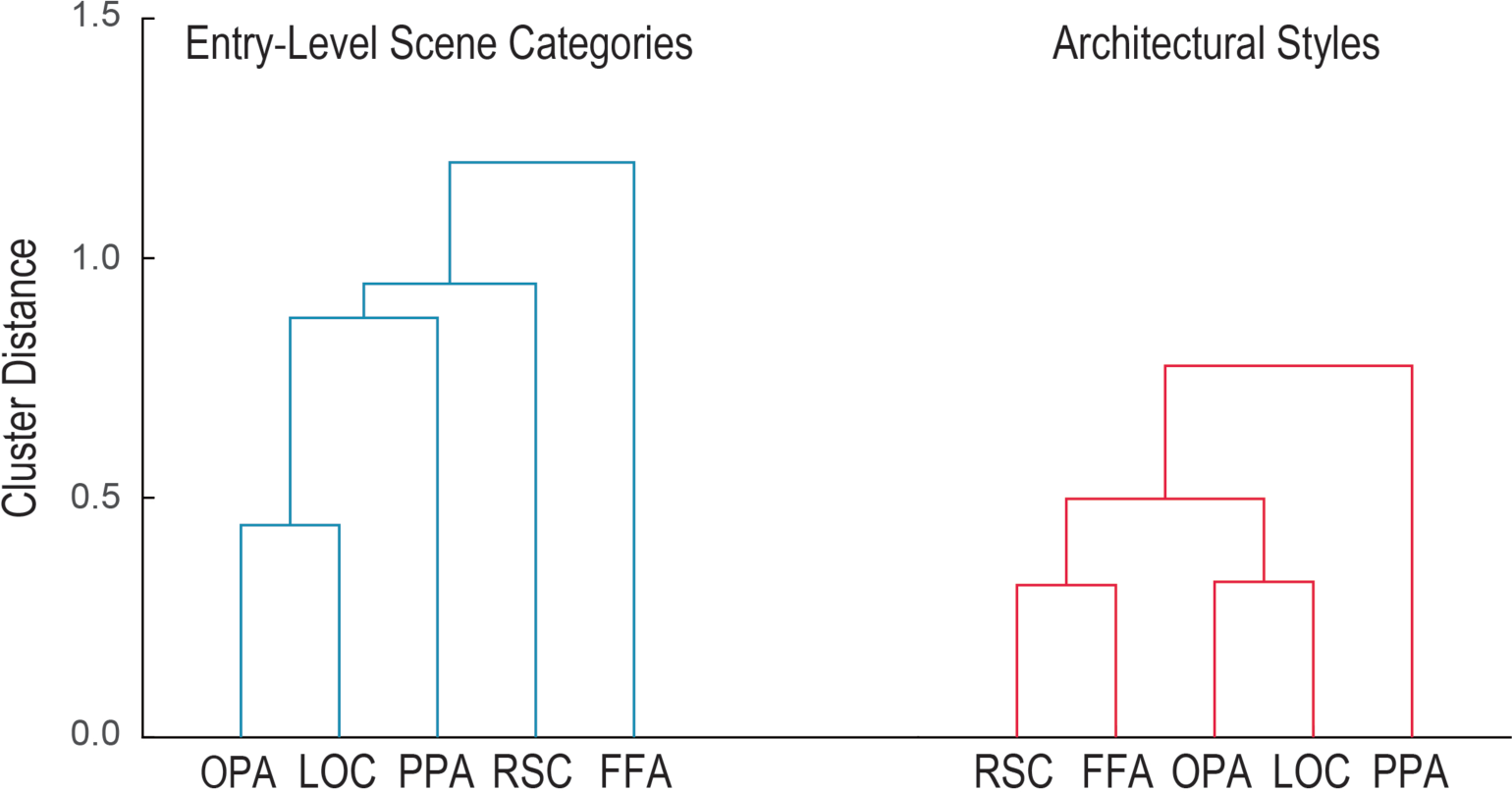
Dendrogram of hierarchical clusters of decoding error patterns from the PPA, OPA, RSC, LOC, and FFA for (A) entry-level scene categories (in blue) and (B) architectural styles (in red). The nearest neighbor linkage method was used to compute cluster distances across the error patterns.

Searchlight analysis of the scanned parts of the brain confirmed the ROI-based results (details of the searchlight analysis are given in the Supplementary Methods). The searchlight map of decoding entry-level scene categories showed significant clusters at both occipital poles and calcarine gyri as well in bilateral lingual, fusiform, and parahippocampal gyri and bilateral transverse occipital sulci. On the other hand, the searchlight map of decoding architectural styles showed clusters encompassing bilateral fusiform gyri and transverse occipital sulci, but not the occipital poles and nearby areas. The searchlight map for decoding buildings by famous architects was similar to that of decoding architectural styles, with an additional small cluster on the left occipital pole. Two significant clusters were found for decoding of facial identity, encompassing parts of occipital cortex and adjacent parietal tissue. Table 1 provides a full list of significant clusters from each searchlight map and Supplementary Figs. S2-5 show significant clusters in axial views separately for the searchlight maps. Analysis of the overlap of individual’s searchlight maps with their ROIs is shown in Supplementary Table S3.

**Table 1.**
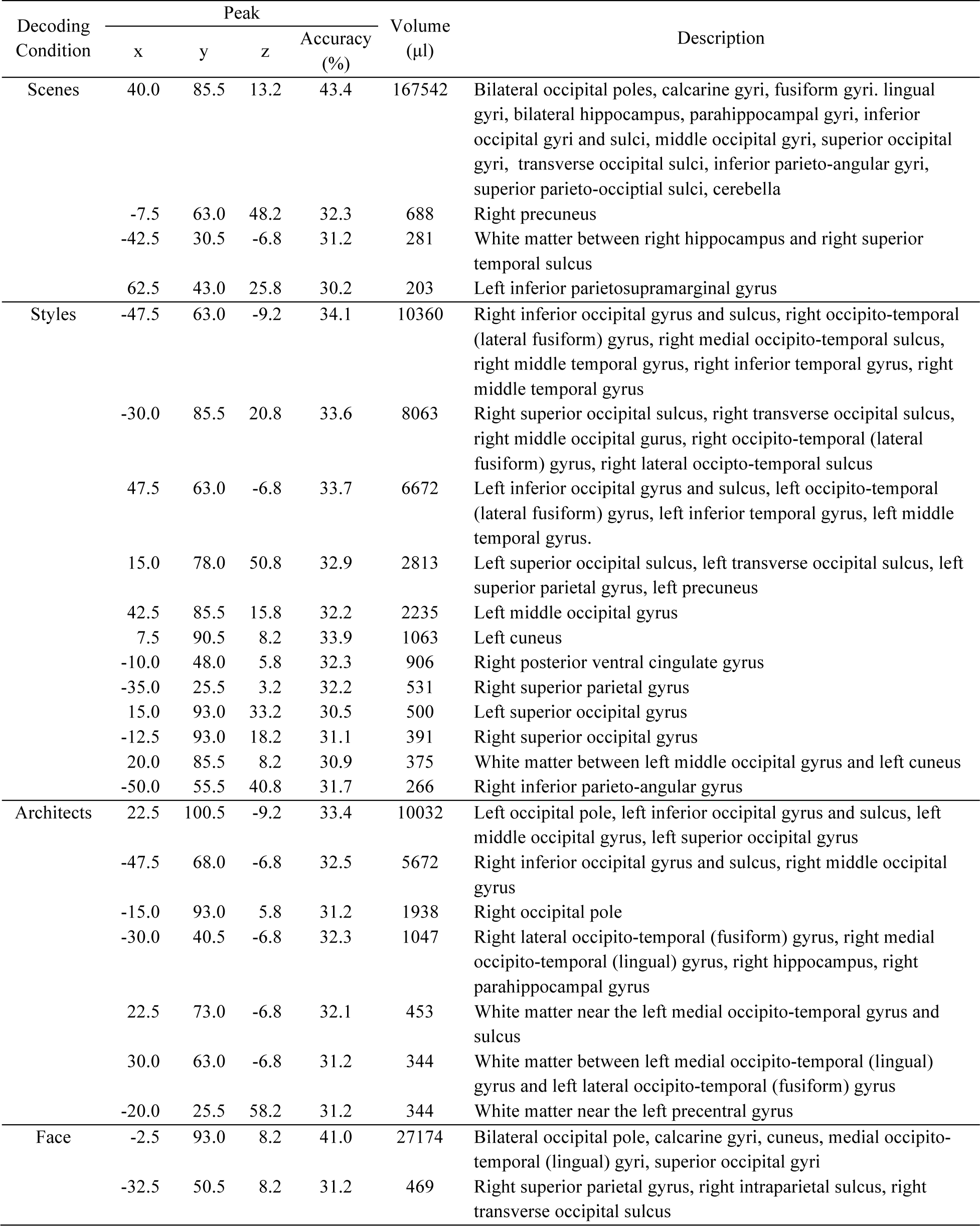
Clusters identified in the searchlight analysis for the four categorization conditions. Significance was determined using *p* < .005 (one-tailed) with a cluster correction (minimum cluster size of 13 voxels).

To explore the nature of the underlying categorical structure of architectural styles in visual cortex in more detail, we analyzed patterns of decoding errors. Decoding errors were recorded in confusion matrices, whose rows (*r*) indicate the ground truth of the presented category, and whose columns (*c*) represent predictions by the decoder. Individual cells (*r,c*) contain the proportion of blocks with category *r*, which were decoded as category *c*. Diagonal elements contain correct predictions, summarized as decoding accuracy in Fig. 2. Off-diagonal elements represent decoding errors. The patterns of decoding errors serve as a proxy for the underlying categorical structure between sub-categories in a particular brain region. We computed the correlations of error patterns as a measure of the similarity between these neural representations across ROIs. Significance of error correlations was established non-parametrically using a permutation test. We also subjected these error correlations to a hierarchical clustering analysis to capture the similarities in categorical structures underlying successful category decoding across high-level visual regions.

In the case of entry-level scene categorization, we found significant correlations of error patterns between the three ROIs known to specialize in scene perception: the PPA, the RSC, and the OPA. We also found significant error correlation between the PPA and the LOC. The FFA did not correlate significantly with any of the other ROIs, even though we could decode entry-level scene categories from the FFA. The hierarchical clustering analysis further illustrates these results by showing a cluster consisting of the OPA and the LOC, which subsequently clustered with the PPA and then the RSC. Note that the error pattern from the FFA was not clustered with any of the other ROIs (Fig. 2A).

For architectural styles, we found a different error correlation structure, showing statistically significant error correlations between the FFA and the high-level visual regions of the PPA, OPA, and LOC. The error correlation between the PPA and the LOC was also significant. Similarly, hierarchical clustering showed that the FFA error pattern was closely clustered with scene-specific brain regions, starting with the RSC error pattern, and subsequently with a cluster consisting of the OPA and the LOC, thus leaving the PPA clustered with the rest of the ROIs (Fig. 2B).

The differences in error pattern similarity structure between entry-level categorization and categorization of architectural styles largely stem from tighter integration of the FFA with the rest of the high-level visual ROIs. In both cases of categorization, error pattern correlations are significant across the scene-specific visual regions – the PPA and OPA as well as the PPA and the LOC. Consistently, cluster distances between the RSC, OPA, LOC, and the PPA are similar between the two experimental conditions. Unlike the case of entry-level scene categorization, the FFA is recruited into the scene processing network for more specialized and demanding subordinate-level scene categorization.

We found low correlations of error patterns between ROIs for decoding architects because of the difficulty of decoding architects from some of the ROIs (i.e., the RSC, the LOC, and the FFA) in the first place. Given that facial identity could not be decoded from any of the high-level visual ROIs, we did not further pursue error correlations for the face identification condition. The decoding error patterns of all four image categories across all six ROIs are shown in Supplementary Fig. S6.

### No discernable effect of expertise in visual cortex

Architectural styles are ultimately visual categories – the majority of defining attributes is visual in nature. Still, accurate recognition of architectural styles or architects of buildings is often affected not only by visual consistency within a style or an architect, but also by the historical, regional, and cultural context of buildings. Prior knowledge of a building’s style may influence the perception of architectural categories and their neural correlates.

Nevertheless, we found no group effect on decoding of either architectural styles or architect, indicating that expertise has little influence on amounts of decodable information of architectural categories in visual cortex – at least not sufficiently to be detected in our experimental paradigm. A mixed ANOVA of decoding accuracy with group as a between-subject factor and visual category as a within-subject factor failed to find a main effect of group or an interaction between group and visual category in any of the ROIs (see Supplementary Table S4). Could these results be due to the lack of the differences in ability to distinguish between architectural styles or architects between the two groups?

To test whether the participants majoring in architecture indeed had higher domain knowledge than the participants majoring in either psychology or neuroscience, we measured expertise for architectural styles in a post-scan behavioral experiment employing the Vanderbilt Expertise Test [8]. In this test, participants were asked to identify which of three displayed images belonged to a given set of six target categories. We confirmed that architecture students had higher expertise for architectural styles and buildings by famous architects: We not only found a significant main effects of group, 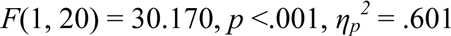, but also a significant interaction between group and visual category, 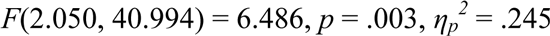. The effect of visual category was also significant, 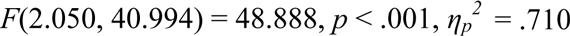. Note, however, that both experts and non-experts could reliably detect target architectural styles from distractors well above chance, as shown in Fig. 3A, consistent with successful decoding of architectural styles in high-level visual regions in both groups.

**Figure 3.**
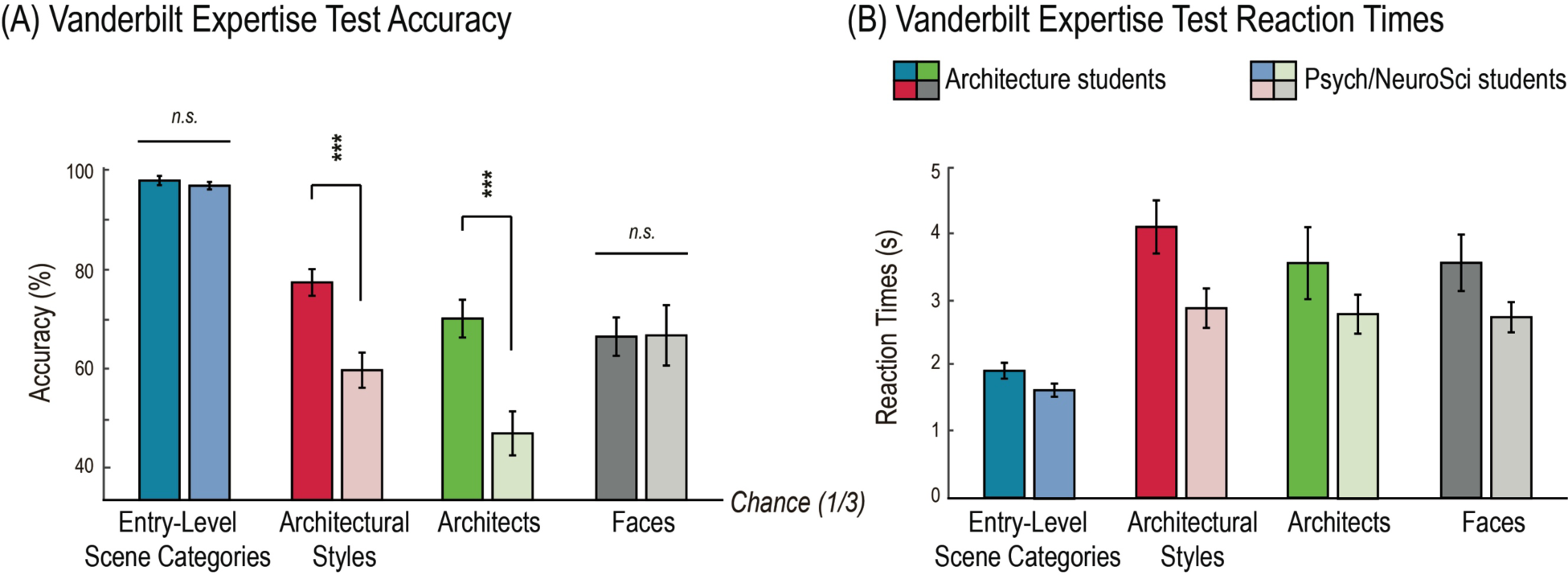
Group-average accuracy rates (A) and reaction times (B) for the four categorization tasks. Dark-colored bars indicate behavioral performances of the eleven architecture students, and bright-colored bars indicate performances of the eleven psychology and neuroscience students. Different hues indicate the four visual categories: entry-level scene categories in blue, architectural styles in red, architects in green, and face identities in gray. Error bars show standard errors of mean. Accuracy of post-hoc comparisons is indicated above the bars. \*\*\**p* <.001.

The same analyses on average reaction times (RT) showed significant main effects of group, 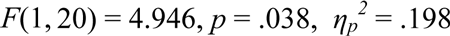, and visual category, 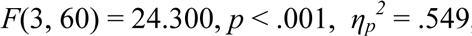, but no significant interaction between them, 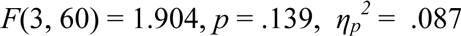, indicating that architecture students (mean RT = 2485 ms) were slower than psychology/neuroscience students (mean RT = 3342 ms) for all types of categorization tasks.

As shown in Fig. 3, post-hoc tests confirmed significantly higher accuracy for students of architecture than for students of psychology/neuroscience for architectural styles, *t*(20) = 3.963, *p* < .001, Cohen’s d = 1.690, and architects, *t*(20) = 6.219, *p* < .001, Cohen’s d = 2.652, but not for entry-level categories, *t*(20) = 1.845, *p* = .080, Cohen’s d = .787, or faces, *t(*20) = .545, *p* = .592, Cohen’s d = .232.

## Discussion

The current study shows for the first time that architectural styles of buildings can be decoded from the neural activity patterns of several high-level visual areas in human temporal cortex. It was even possible to decode the architects of buildings from neural activity elicited by images of the buildings in the PPA and the OPA. However, architectural styles, unlike entry-level scene categories, could not be decoded from V1, indicating that the simple visual properties encoded in V1 are insufficient to discriminate between architectural styles. We also found substantial similarity in error patterns of decoding architectural styles between the FFA and the other highlevel visual regions.

The PPA is one of the most robust modular regions known to be specialized for outdoor and indoor scenes and buildings [9]. Thus, it is no surprise to find a significant amount of decodable information about architectural styles in this “building” area. How exactly the PPA encodes various scenes and buildings, however, is not entirely clear, since the PPA contains decodable information about numerous aspects of scenes, such as spatial structure [5, 10, 11], texture and material properties [12], as well as semantic categories [6, 7]. Note that all these perceptual aspects are critical for characterizing architectural styles. The PPA is also suggested as a key area for linking various perceptual instantiations of the same building [13], and similar neural mechanisms may be at work to form subordinate-level categories of buildings, such as for architectural styles. We suggest that the PPA elicits style-specific neural activity patterns for buildings by extracting multi-dimensional statistics specific to architectural styles from a given instance.

Architectural styles of buildings could also be decoded from activity patterns in the OPA and RSC, the other two scene-specific ROIs [14, 15]. The OPA has been suggested to contain primitive scene representations by encoding mid-level visual properties [14, 16]. Since we could not decode architectural styles from V1, it is likely that the neural representations of architectural features begin to arise from mid-level visual features (e.g., symmetry, curvature, collinearity etc.), which are available in the OPA. Mid-level visual properties, in fact, have previously been suggested to contribute to successful cross-decoding between interior and exterior views of landmark buildings in the OPA [13].

The RSC, on the other hand, has been found to reflect mnemonic and contextual aspects of realworld scenes rather than their perceptual aspects [15, 17]. The building images were all relatively famous landmarks, and might have induced semantic or even episodic memory components. Given our data, however, we cannot provide an operational mechanism of how the RSC can differentially respond to buildings according to their architectural styles.

Real-world scene categories elicit distributed neural activity patterns in visual regions sensitive to objects. Scene information in the LOC has been associated with category-specific object statistics [10, 16, 18] – office scenes are highly likely to contain desks, chairs, and computers whereas city street scenes are highly likely to contain vehicles, buildings, and driveways. Despite the high homogeneity in object statistics (i.e., buildings and occasionally trees and vehicles), we still found successful decoding of architectural styles from the LOC as well as a significant error pattern correlation between the LOC and the PPA. We, therefore, suggest that the contribution of the LOC may be to encode local elements [19, 20] such as motifs and embellishments common in an architectural style. However, this conjecture will have to be tested rigorously in future investigations.

Surprisingly, architectural styles also can be decoded from another high-level visual region typically considered as not preferring scenes and buildings – the fusiform face area, previously implicated in the preferential processing of faces [21] as well as visual expertise [22]. More importantly, the FFA is recruited as part of a network of regions that share similar error patterns. By contrast, entry-level categorization of scenes does not include the FFA in the same way, instead relying on a tight network of three scene-selective areas, the PPA, the RSC, and the OPA, as well as the LOC. The FFA could be involved in the encoding of configural characteristics of buildings. This is consistent with the FFA’s role in visual expertise as shown for object categories as varied as birds, cars, motorcycles or artificial “Greeble” objects [22]. Note that those results were shown for mean activity levels, whereas ours appear in the interpretation of multi-voxel patterns of brain activity.

Taken together, the hallmark of neural representations of architectural styles is their distributed and interactive nature. It may be, in fact, the only practical solution for human visual cortex to deal with the multi-dimensionality underlying the visual classification between architectural styles. To characterize a style, one needs to inspect global shape and layout of a building, the shape of architectural elements (i.e., roof, walls, pillars) and their configurations, construction materials, local motif and embellishments, etc. For instance, byzantine architecture is characterized by symmetry in the global shape of buildings and a dome roof, stone brick exterior, and tile mosaic embellishments, whereas deconstructive architecture is well known for its noncollinearity and fragmented global shape and concrete, steel, glass exterior, and minimal embellishments. These insights may in fact explain the lack of decodable information about architectural styles in V1: the multi-dimensionality of distinguishing features may be simply beyond the processing capability of V1. Similarly, the neural representations of artistic styles associated with painters (Dali or Picasso) have been found to reflect painter-specific global visual statistics (i.e., chromatic intensity histograms) rather than pixel-based features [23].

Unexpectedly, we did not find a reliable effect of domain expertise on the amount of decodable information of architectural styles in the visual cortex. Categorizing a building by its architectural style or its designer involves not only detecting characteristic visual features, but also recruitment of semantic knowledge. A number of past studies suggest that gaining expertise of perceptual categorization relies on intercortical loops involving not only modality-specific sensory regions (i.e. visual cortex), but also medial temporal, parietal, and prefrontal cortex and subcortical structures such as basal ganglia (for a review, see [24]). Consistent with this idea, we also found decodable information about architectural features in parietal regions beyond visual cortex. Another possibility is that core differences between experts and non-experts prevail in their post-perceptual analyses of buildings involving cognitive and aesthetic appreciation [3, 25] rather than perceptual analyses.

In summary, several high-level visual regions, but not primary visual cortex, contain decodable neural representations of architectural styles and architects of buildings. The FFA substantially participates in a network of high-level visual areas characterized by similar error patterns in the decoding architectural styles but not in decoding entry-level scene categories. We showed that high-level visual regions in the human brain contain neural correlates of the visual perception of architectural styles, which are likely to be driven by complex perceptual statistics specific to an architectural style or an architect. We found no evidence for differences in the neural code in visual cortex between experts and non-experts for architecture. Our findings have characterized neural mechanisms for perceptual encoding of architecture in the human visual system, one of the predominant and longest-lasting artefacts of human culture.

## Methods

All experimental procedures were approved by the institutional review board of The Ohio State University, and all data collection and analyses were carried out in accordance with the approved guidelines.

### Participants

Twenty-three healthy undergraduate students in their final year at The Ohio State University participated in the study for monetary compensation of $15/hour and gave written informed consent. We recruited eleven students from the Department of Architecture (2 females; l left-handed; age range = 21–27, *M* = 22.4, *SD* = 3.0), and twelve senior students majoring in psychology or neuroscience (3 females; 2 left-handed, age range = 21– 24, *M* = 21.8, *SD* = 0.9). Data from one psychology student was not included in the analysis due to excessive head motion during the scan.

### Post-scan behavioral experiment

We measured participants’ visual domain knowledge in a post-scan behavioral experiment similar to the Vanderbilt Expertise Test [8]. Domain knowledge for each visual category was tested in four separate blocks. Each block consisted of three components: study, practice, and testing. During study, participants were introduced to six target categories. Example images for each of the six target categories were displayed on the screen with correct category labels: (1) architectural styles: Byzantine, Gothic, Renaissance, Modern, Postmodern, and Deconstructive; (2) buildings by famous architects: Peter Eisenman, Antoni Gaudi, Frank Gehry, Michael Graves, Le Corbusier, and Frank Lloyd-Wright; (3) entry-level scene categories: fountains, highways, mountains, pastures, skylines, and waterfalls; (4) faces: six non-famous individuals varied in gender and race. Following the study phase, participants experienced twelve practice trials. In these trials, three images (12° x 12° of visual angle each) were presented side by side. Participants were asked to indicate which of the three images belonged to a given target category by pressing one of the keys, “1,” “2,” or “3.” During practice, one of the three images was always drawn from the set of studied examples. The images were presented until the participant made a response, and feedback was provided by displaying the word “CORRECT” or “INCORRECT.” Study exemplars were shown again halfway through practice and at the beginning of the subsequent test phase. For the 35 test trials, 24 new grayscale images from the target categories and 48 new grayscale foil images from different categories were used. Structure of the test trials was the same as practice, except that participants no longer received feedback. The entire experiment lasted approximately 30 min.

The average accuracy rates and reaction times of individual participants entered an ANOVA using participant group as a between-subjects factor and visual category (entry-level scene categories vs. architectural styles vs. architects vs. faces) as a within-subjects factor. The degrees of freedom were adjusted using Greenhouse-Geisser, because the assumption of equal variance was violated for the factor of visual category. To further analyze the significant group x visual category interaction, post-hoc t-tests were performed, using Bonferroni correction to account for multiple comparisons (critical p = .0125).

### fMRI Experiment

MRI images were recorded on a 3T Siemens MAGNETOM Trio MRI scanner with a 12-channel head coil at the Center for Cognitive and Behavioral Brain Imaging at The Ohio State University. High-resolution anatomical images were obtained with a 3D-MPRAGE sequence with coronal slices covering the whole brain; inversion time = 930 ms, repetition time (TR) = 1900 ms, echo time (TE) = 4.44 ms, flip angle = 9°, voxel size = 1 x 1 x 1 mm, matrix size = 224 x 256 x 160. Functional images were obtained with T2*-weighted echo-planar sequences with coronal slices covering approximately the posterior 70% of the brain: TR = 2000ms, TE = 28ms, flip angle = 72°, voxel size = 2.5 x 2.5 x 2.5 mm, matrix size = 90 x 100 x 35.

Participants viewed 512 grayscale photographs of four visual categories: (1) 32 images of representative buildings of each of four architectural styles: Byzantine, Renaissance, Modern, and Deconstructive; (2) 32 images of buildings designed by each of four well-known architects: Le Corbusier, Antoni Gaudi, Frank Gehry, and Frank Lloyd-Wright; (3) 32 scene images per each of four entry-level scene categories: mountains, pastures, highways, and playgrounds; (4) 32 face images per each of four different individuals [26]. The building images encompassed a variety of views, including close-ups, far views, and aerial views. This variation of views ensured that building categories were not confounded with other global scene properties, such as openness or mean distances [5, 10, 27]. Brightness and contrast were equalized across all images. Images were back-projected with a DLP projector (Christie DS+6K-M 3-chip SXGA+) onto a screen mounted in the back of the scanner bore and viewed through a mirror attached to the head coil. Images subtended approximately 12º x 12 º of visual angle. A fixation cross measuring 0.5º x 0.5º of visual angle was displayed at the center of the screen.

During each of nine runs, participants saw sixteen 8-second blocks of images. In each block, four photographs from a single category were each shown for 1800 ms, followed by a 200 ms gap. The order of images within a block and the order of blocks within a run were randomized in such a way that the four blocks belonging to the same stimulus type (entry-level scenes, styles, architects, faces) were shown back to back. A 12-sec fixation period was placed between blocks as well as at the beginning and the end of each run, resulting in a duration of 5 min 32sec per run. Occasionally, (approximately one out of eight blocks), an image was repeated back-to-back within a block. Participants were asked to press a button when they detected image repetitions.

FMRI data were motion corrected, spatially smoothed (2 mm full width at half maximum), and converted to percent signal change. We used a general linear model with only nuisance regressors to regress out effects of motion and scanner drift. Residuals corresponding to image blocks were extracted with a 4 s hemodynamic lag and averaged over the duration of each block. Block-average activity patterns within pre-defined ROIs was used for multi-voxel pattern analysis (MVPA).

MVPA was performed separately for each participant by training a linear support vector machine on the data for all runs except one, and then testing on the data from the left-out run. In a leaveone-run-out (LORO) cross validation each run was left out in turn, thus generating predictions for each run. Separate decoders were trained for each participant, each ROI, and each visual category (entry-level scene categories, architectural styles, buildings by famous architects, faces). Proportion of correct predictions is reported as accuracy. At the group level, accuracy is compared to chance (0.25) using one-tailed t tests. P-values were adjusted using false discovery rate [28].

Prediction errors were recorded in a confusion matrix. Patterns of errors (off-diagonal elements of the confusion matrices) were correlated between ROIs. Significance of error correlations was tested non-parametrically against the null distribution of correlations obtained by jointly permuting the rows and columns of one of the confusion matrices. Only error correlations with none of the 24 permutations resulting in higher correlation than the correct ordering were deemed significant. Correlations of group-level error patterns between ROIs form the basis of a hierarchical clustering analysis, performed in MATLAB with a nearest-neighbor linkage method to illustrate the error pattern correlations across the ROIs as dendrograms.

## Author Contributions

Conceptualization, D. B. W, J. N., B. N., and H. C.; Methodology, H. C., D. B. W., B. N.; Investigation, H. C., and D. B. W.; Writing – Original Draft, H. C.; Review & Editing, D. B. W., H.C., and J. N.; Funding Acquisition, D. B. W.; Resources, D. B. W.; Supervision, D. B. W. and J. N.

## Additional Information

*Competing financial interests*: The authors declare no competing financial interests.

